# *Clostridioides difficile* infection increases circulating p-cresol levels and dysregulates brain dopamine metabolism: linking gut-brain axis to autism spectrum disorders?

**DOI:** 10.1101/2021.10.22.465382

**Authors:** Akhil A. Vinithakumari, Piyush Padhi, Belen Hernandez, Susanne Je-Han Lin, Aaron Dunkerson-Kurzhumov, Lucas Showman, Matthew Breitzman, Caroline Stokes, Yousuf Sulaiman, Chandra Tangudu, Deepa Ashwarya Kuttappan, Muhammed Shafeekh Muyyarikkandy, Gregory Phillips, Vellareddy Anantharam, Ann Perera, Brett Sponseller, Anumantha Kanthasamy, Shankumar Mooyottu

## Abstract

Gastrointestinal illnesses are one of the most common comorbidities reported in patients with neurodevelopmental diseases, including autism spectrum disorders (ASD). Gut dysbiosis, overgrowth of *C. difficile*, and gut microbiota-associated alterations in central neurotransmission have been implicated in ASD, where the dopaminergic axis plays an important role in the disease pathogenesis. Human *C. difficile* strains produce a significant amount of the toxic metabolite p-cresol, an inhibitor of dopamine beta-hydroxylase (DBH), which catalyzes the conversion of dopamine (DA) to norepinephrine (NE). p-Cresol is known to precipitate and exacerbate autistic behavior in rodents by increasing DA levels and altering DA receptor sensitivity in brain regions relevant to ASD. Therefore, we hypothesized that *C. difficile* infection dysregulates dopaminergic metabolism by increasing p-cresol levels in the gut and systemic circulation, and by inhibiting brain DBH, ultimately leading to elevated DA in different brain regions. For testing this hypothesis, we induced antibiotic-associated *C. difficile* infection in mice and determined the gut and serum p-cresol levels, serum DBH activity, and dopamine and its metabolite levels in different brain regions relevant to ASD. The results showed that *C. difficile* infection causes a significant increase in striatal DA, accompanied by significantly altered levels of DA metabolites and NE in different brain regions (p < 0.05). In addition, significantly increased circulating p-cresol levels and reduced DBH activity were observed in *C. difficile* infected mice (p < 0.05). Therefore, the results from this study suggest a potential link between *C. difficile* infection and alterations in the dopaminergic axis implicated in the precipitation and aggravation of ASD.

## Introduction

Although the involvement of the gut-brain axis is increasingly implicated in the precipitation and manifestation of neurodevelopmental disorders such as autism, mechanistic evidence that directly connects gut bacteria, bacterial metabolites, and neuronal dysregulation is sparse.^1, 2^ Disruption of the normal gut microbiota is known to cause an overgrowth of unfavorable bacterial communities that increase toxic metabolites in the gut.^3^ Several of these metabolites are known to impact neurotransmission and brain functions, underscoring the importance of the gut-brain axis in health and disease.^4^

p-Cresol is a toxic phenolic compound produced by bacterial fermentation of protein in the human large intestine.^5^ p-Cresol is readily absorbed into the bloodstream and secreted in urine.^6, 7^ Moreover, it is already known that patients with autism spectrum disorders (ASD) often have a significantly higher urine p-cresol level, which is considered a biomarker for this disease complex.^8–10^ A recent study demonstrated that intravenous p-cresol injection aggravates autistic behaviors in a murine genetic model of ASD by dysregulating the dopaminergic axis.^11^ Another study revealed ASD-like behavior in rats intraperitoneally administered with p-cresol.^12^ Similar results with dysregulation in dopaminergic neurotransmission were demonstrated following oral p-cresol administration in non-mutant mice.^13^ Therefore, p-cresol mediated alterations in the dopaminergic pathway appear to be associated with the pathophysiology of ASD.

p-Cresol is a potent inhibitor of the enzyme dopamine beta-hydroxylase (DBH) that converts dopamine (DA) to norepinephrine (NE).^14–16^ Thus, inhibition of DBH increases the concentration of DA in the central nervous system. Notably, a low maternal serum level of DBH and mutations in the DBH genes have been associated with autism in children.^17, 18^ Additionally, increased urinary and cerebrospinal fluid levels of homovanillic acid, a DA metabolite, have been reported in autistic patients.^19–21^ Moreover, an elevated brain DA level, mutations in the DA transporter protein, altered sensitivity of DA receptors, and impaired striatal dopaminergic neurotransmission are observed in autistic patients and animal models of ASD.^22–25^ In addition, DA receptor antagonists are found to reduce autistic symptoms in patients.^25, 26^

Gut dysbiosis induced by prolonged antibiotic therapy is the most common cause of *C. difficile* overgrowth and associated diarrhea in infants and adults.^27, 28^ Human *C. difficile* strains produce a significant amount of p-cresol, unlike most other gut bacteria.^29–31^ *C. difficile* produces 10 to 1000 times more p-cresol than other known p-cresol producing bacteria in the gut.^5, 32^ Indeed, recent studies demonstrated p-cresol induced autistic behavior in mice by altering the dopaminergic axis.^11, 13^ What remains unknown specifically is the p-cresol mediated functional consequence of *C. difficile* infection on the dopaminergic neurotransmission from a gut-brain axis standpoint. Moreover, the involvement of possible mediators, such as DBH, that directly connects gut microbial p-cresol to DA dysmetabolism needs to be demonstrated. Thus, the current work investigates the effect of antibiotic-induced *C. difficile* infection and consequent *in vivo* production of p-cresol on the DA metabolism in the brain.

## Results

### C. difficile UK1 produces a significant amount of p-cresol in vitro

To determine the p-cresol producing ability of the human hypervirulent *C. difficile* strain (UK1), we performed a p-HPA-p-cresol conversion assay. For this experiment, we quantified concentrations of total p-cresol and its substrate para-hydroxyphenyl acetic acid (p-HPA) concentrations at different time points in *C. difficile* UK1 cultures by high-performance liquid chromatography (HPLC) *in vitro*. Our results indicated that *C. difficile* UK1 completely metabolized p-HPA to p-cresol in brain heart infusion (BHI) media by 24h of incubation in anaerobic conditions **(Figure 1)**. This data confirm the p-cresol producing capability of human hypervirulent *C. difficile* strain, suggesting that this bacterium could increase the gut p-cresol concentrations in *C. difficile* infected host.

**Figure 1.**
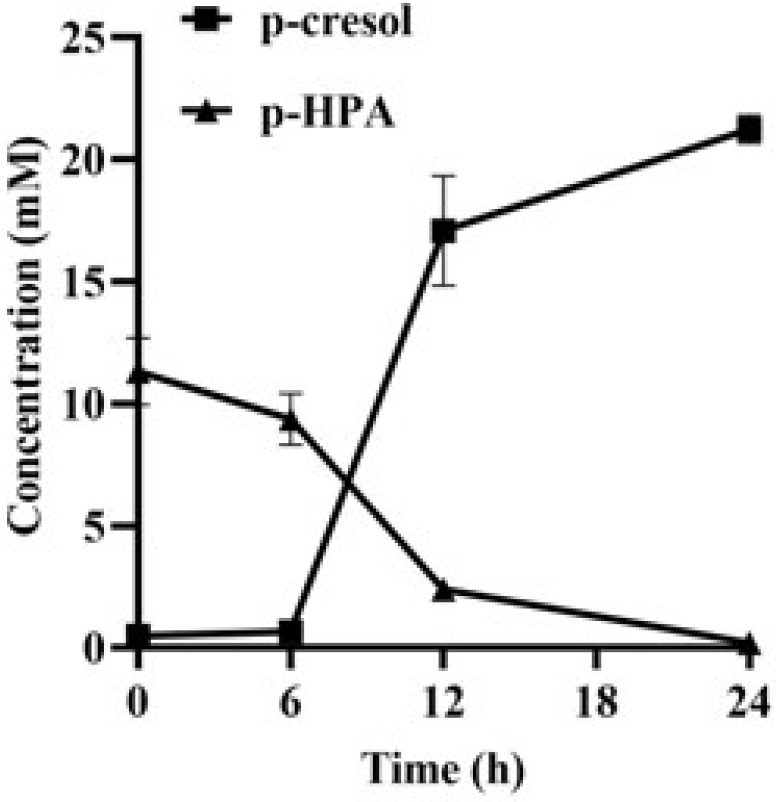
*C. difficile* UK1 produces a significant amount of p-cresol in vitro: *C. difficile* was cultured anaerobically in supplemented brain heart infusion (BHIS) broth with the substrate 0.3 % p-HPA, and the concentrations of p-HPA and newly formed p-cresol in culture supernatant at different time points were quantified using HPLC analysis. *C. difficile* metabolized the substrate p-HPA within 24h.

### Antibiotic induced C. difficile infection increases gut p-cresol concentration

In order to quantitate *in vivo C. difficile* p-cresol production in the murine gut, we performed a *C. difficile* challenge study. For this experiment, 3-week-old C57BL/6 mice were administered with an oral antibiotic cocktail followed by an intraperitoneal clindamycin injection in order to induce gut dysbiosis. Gut dysbiosis is required for *C. difficile* colonization in mice as in humans.^33, 34^ Mice were then challenged with 10^4^ *C. difficile* UK1 spores **(Figure 2A)**. From the previous experiments conducted by the authors and other investigators, it is known that 10^4^ *C. difficile* UK1 spores reliably induces clinical CDI with less mortality in mice compared to high dose (10^6^ to 10^7^ spores) challenge, simulating chronic persistent CDI in humans. Diarrhea or wet tail were noticed in all *C. difficile* challenged mice on day-2 post-inoculation. Histopathologic examination of the colonic sections revealed no significant lesions in the colon of the control and antibiotic-control (Abx) groups **(Figure 2B)**. As expected, the colon of the mice challenged with 10^4^ *C. difficile* UK1 (C.Diff) exhibited significant pathologic changes in the mucosal epithelium, as evidenced by necrosis, moderate neutrophilic inflammatory infiltrates in the lamina propria and submucosa and marked submucosal edema, contributing to a significantly increased cumulative colitis score **(Figure 2B & 2C)**. In addition, the total cecal p-cresol level (free and bound) on day-2 post-challenge was determined using HPLC method **(Figure 2D)**.^35^ A significant increase in total cecal p-cresol concentration was observed in C. Diff group compared to controls (p < 0.05). An increasing trend in the cecal p-cresol, although not statistically significant was observed in Abx group compared to control **(Figure 2D)**. Thus, the data indicate that clinical *C. difficile* infection significantly increases gut p-cresol levels in infected mice.

**Figure 2.**
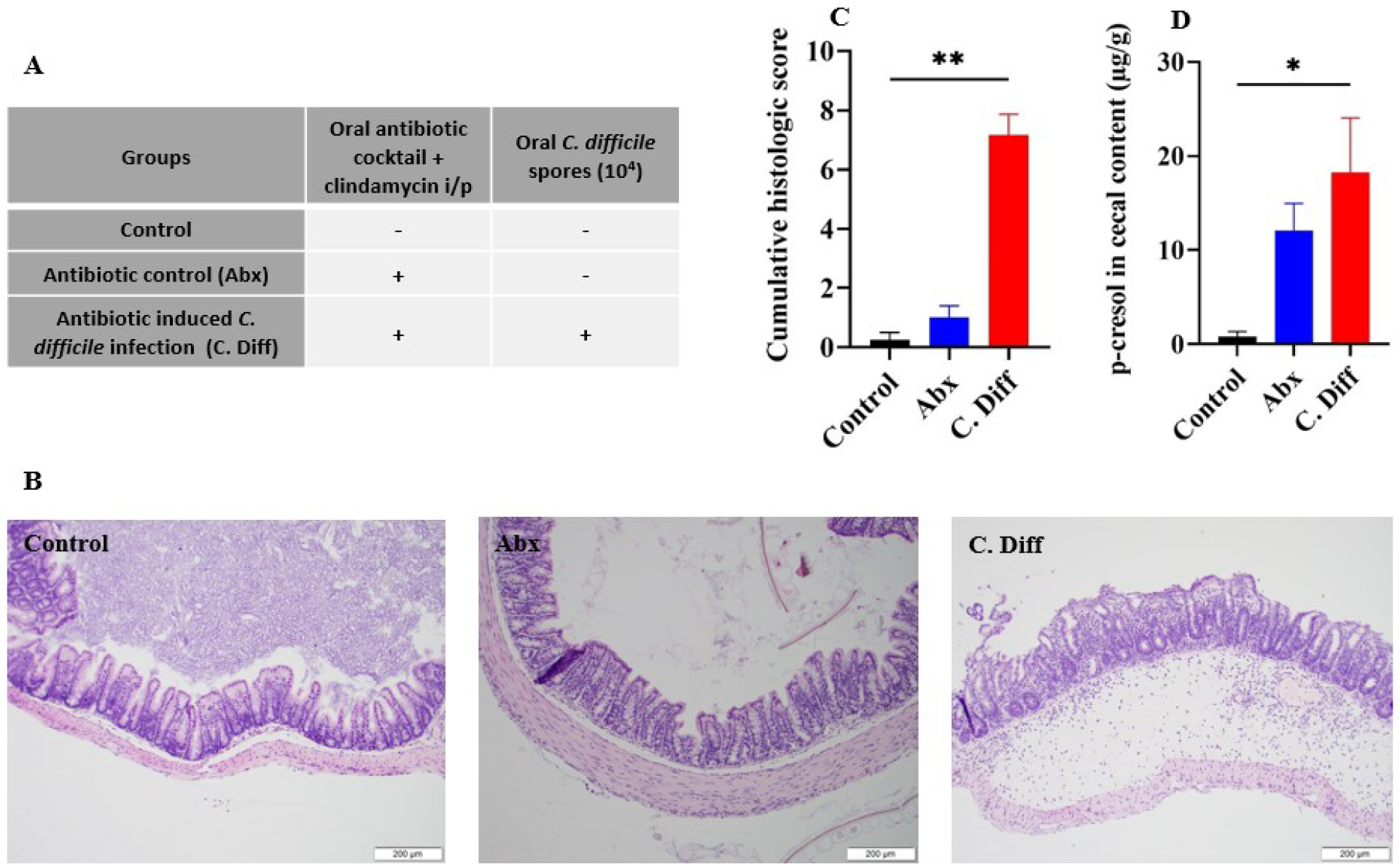
Antibiotic induced *C. difficile* infection increases gut p-cresol concentration: **(A)** *Experimental outline:* Three-four weeks old C57BL/6 mice are treated with an oral antibiotic cocktail or PBS and an intraperitoneal clindamycin injection or PBS for inducing gut dysbiosis and then challenged with 10^4^ *C. difficile* spores or PBS. Animals were euthanized, and cecal tissues were collected approximately two days post-*C. difficile* challenge; **(B)** *Histologic lesions in different treatment groups:* mice colons from each treatment group were processed and stained with hematoxylin and eosin. Minimal edema, minimal to absent neutrophilic infiltration, and no necrosis were observed in the colonic epithelium of the Control and Abx groups. Significant epithelial necrosis, mucosa, and submucosal neutrophilic infiltration and submucosal edema were observed in the colons of the C. Diff group; **(C)** *Histopathology and colitis score:* Histologic scores were assessed by a blinded pathologist across the groups based on epithelial damage, mucosal edema, and neutrophil infiltration. The scores are combined into a cumulative histologic score for each mouse ranging from 0 to 9; **(D)** *Gut p-cresol concentration:* Total cecal p-cresol concentrations in control, antibiotic control (Abx), and *C. difficile* challenge (C. Diff) groups at 2-day post-inoculation, quantitated by HPLC method; **p< 0.01, *p < 0.05.

### C. difficile infection increases serum p-cresol and reduces serum DBH activity

Since p-cresol produced in the intestine is readily absorbed into the blood stream, we investigated the changes in the serum p-cresol levels in *C. difficile* challenged and control groups. Previous studies demonstrated that p -cresol is a neurotoxin that can cross the blood brain barrier, impair neurotransmission, and induce behavioral abnormalities.^12^ In addition, p-cresol is a potent inhibitor of DBH enzyme.^14–16^ It is also known that the serum DBH is predominantly contributed by the brain tissue, and hence the serum DBH activity meaningfully reflects the brain DBH activity.^36, 37^ Therefore, we quantitated serum p-cresol levels in *C. difficile* challenged and control mice. Furthermore, we performed a DBH activity assay to determine whether *C. difficile* induced changes in the circulating p-cresol inhibits the activity of this enzyme, which may reduce DA to NE conversion in brain. As expected, a significant increase in total serum p-cresol concentration was observed in C. Diff group compared to controls (p < 0.05). Minor increases in the serum p-cresol (not significant) were observed in Abx group compared to control **(Figure 3A).** Most importantly, a significant reduction in the serum DBH activity (p < 0.05) was observed in *C. difficile* challenged mice as indicated by the lower response in LC-MS detection of the reaction product (octopamine) compared to controls. **(Figure 3B)**. Thus, the results from this set of experiments demonstrates a significant increase in serum p-cresol concentration and diminished DBH activity in *C. difficile* infected mice.

**Figure 3.**
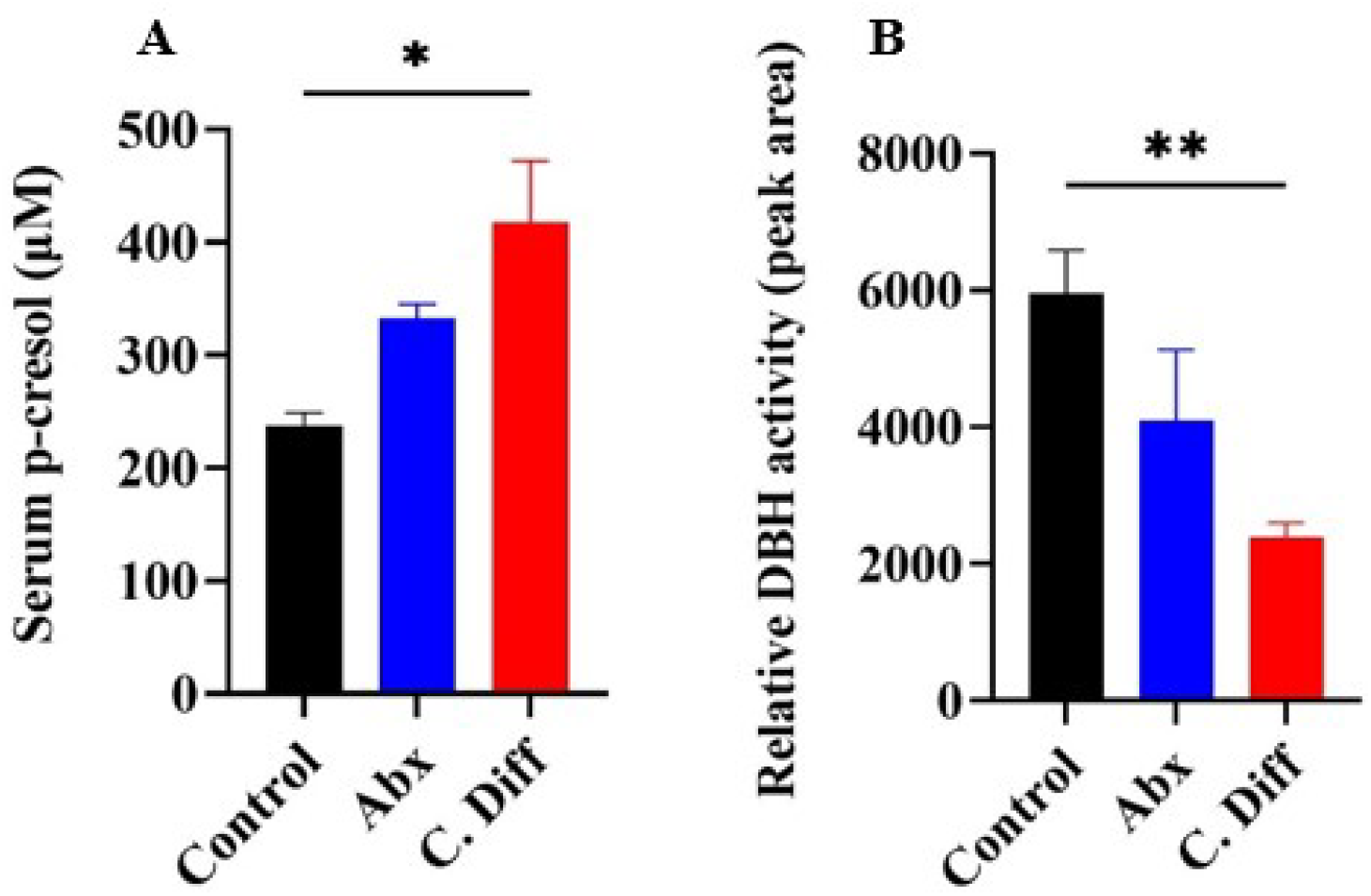
*C. difficile* infection increases serum p-cresol and reduces serum DBH enzyme activity: Three-four weeks old C57BL/6 mice are treated with an oral antibiotic cocktail or PBS and an intraperitoneal clindamycin injection or PBS for inducing gut dysbiosis and then challenged with 10^4^ *C. difficile* spores or PBS. **(A)** *Serum p-cresol:* Sera were collected 2-day post-inoculation to quantify total serum p-cresol concentration in Control, antibiotic control (Abx), and *C. difficile* challenge (C. Diff) groups using the HPLC method**. (B)** *Relative serum DBH activity:* DBH enzyme activity in each treatment group was determined using an enzymatic assay coupled with LC-MS. The relative DBH activity was expressed as the peak area representing the tyramine octopamine transformation per hour in each treatment group. **p< 0.01, *p < 0.05.

### C. difficile infection increased DA and its metabolite levels in corpus striatum and prefrontal cortex

Increased gut and serum concentrations of p-cresol have been demonstrated to alter dopaminergic neurotransmission, which has been implicated in neurological disorders, including autism.^11–13^ Two major dopaminergic pathways in brain significant to ASD include the mesocorticolimbic (MCL) circuit, and the nigrostriatal (NS) circuit.^38, 39^ Substantia nigra and the ventral tegmental area are the two major DA producing areas in the brain. Neurons from the substantia nigra project to the dorsal striatum, forming the nigrostriatal (NS) circuit, which is associated with the motor aspects of goal-directed behavior. The ventral tegmental area projects to the prefrontal cortex and the ventral striatum forming the mesocorticolimbic (MCL) circuit, which is involved in reward and motivation-related behavior. Dysregulation of the mesocorticolimbic circuit leads to social deficits, and malfunction of the nigrostriatal circuit leads to stereotyped behaviors seen in ASD.^39^ Striatum is a critical area that receives projections from both systems; and therefore, DA and its metabolite concentrations were examined in the striatum in response to antibiotic treatment and *C. difficile* infection.

The results from the neurochemical analysis for striatal catecholamines **(Figure 4)** showed a significantly increased DA concentration in the C.Diff group compared to the untreated control (p < 0.05). However, concentrations of DA metabolic products 3,4-dihydroxyphenylacetic acid (DOPAC) and homovanillic acid (HVA) did not significantly differ from that of control animals. Similarly, no significant difference was observed in the NE concentration in the striatum of *C. difficile* challenged mice compared to the negative control. DA concentrations were unaltered or slightly reduced in the antibiotic control group (Abx) compared to the untreated control. In contrast, the significantly lower DOPAC and HVA concentrations (p < 0.05) were observed in Abx group compared to the negative control. No statistically significant difference was observed in the NE levels in Abx compared to the negative control, although a decreasing trend was noted.

**Figure 4.**
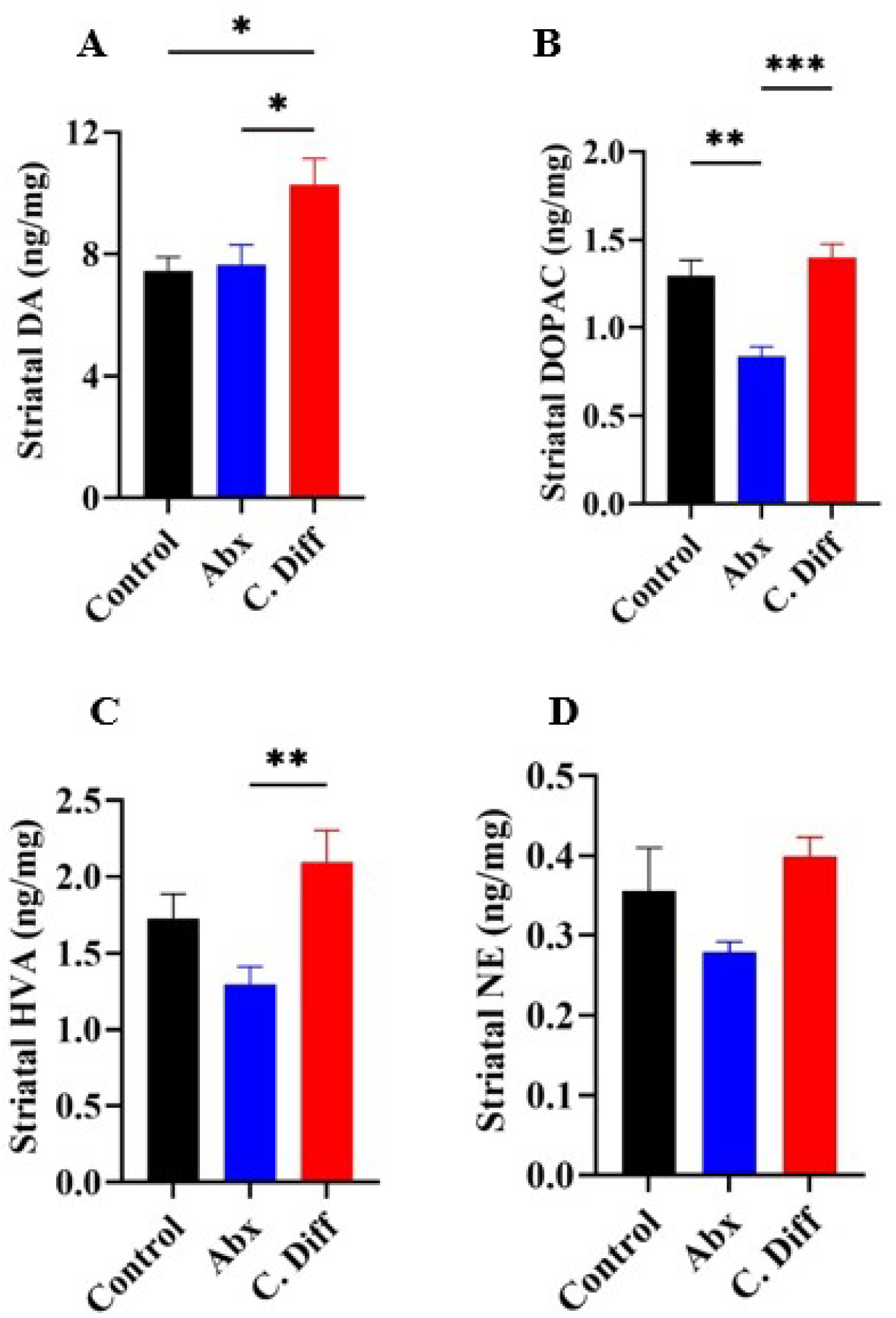
Antibiotic induced *C. difficile* infection increased striatal dopamine and its metabolite levels: Three-four weeks old C57BL/6 mice are treated with an oral antibiotic cocktail or PBS and an intraperitoneal clindamycin injection or PBS for inducing gut dysbiosis and then challenged with 10^4^ *C. difficile* spores or PBS. Striatal catecholamines- **(A)** dopamine (DA), **(B)** its metabolite 3, 4-dihydroxyphenylacetic acid (DOPAC), **(C)** homovanillic acid (HVA), and **(D)** norepinephrine (NE) levels in Control, antibiotic control (Abx), and *C. difficile* challenge (C. Diff) groups at 2-day post-inoculation were quantitated by HPLC-ED method. *p < 0.05

A comparable, similar trend was observed in the catecholamine concentrations in the pre-frontal cortex **(Figure 5)**. An increasing trend in DA and DOPAC concentrations and a statistically significant (p < 0.05) increase in HVA in the *C. difficile* challenged mice were observed when compared to both negative and antibiotic control groups. Interestingly, although statistically not significant, a decreasing trend of NE concentration was observed in C.Diff group compared to both control groups.

**Figure 5.**
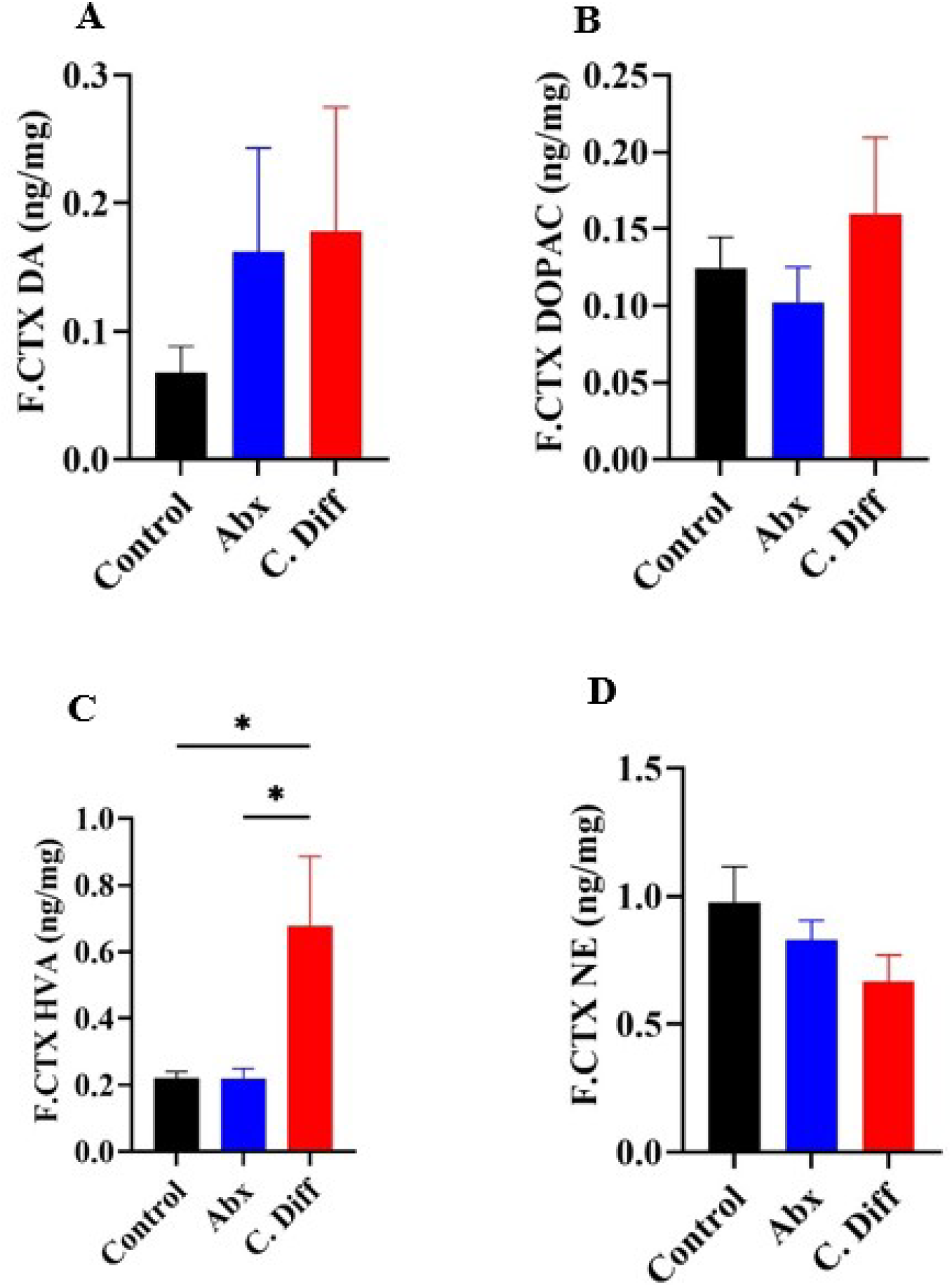
*Antibiotic-induced* C. difficile infection altered catecholamine levels in the prefrontal cortex: Three-four weeks old C57BL/6 mice are treated with an oral antibiotic cocktail or PBS and an intraperitoneal clindamycin injection or PBS for inducing gut dysbiosis and then challenged with 10^4^ *C. difficile* spores or PBS. Prefrontal Cortex (F.CTX) catecholamines- **(A)** dopamine (DA), **(B)** its metabolite 3,4-dihydroxyphenylacetic acid (DOPAC), **(C)** homovanillic acid (HVA), and **(D)** norepinephrine (NE) levels in Control, antibiotic control (Abx), and *C. difficile* challenge (C. Diff) groups at 2-day post-inoculation were quantitated by HPLC-ED method. *p < 0.05.

### Both antibiotic treatment and antibiotic induced *C. difficile* infection decreased NE levels in hippocampus

In contrast to the striatal and prefrontal cortex, the neurochemical profile of catecholamines in the hippocampus revealed significant reduction of the DA degradation product DOPAC (p<0.05) in C.Diff group, accompanied with a statistically nonsignificant, but decreasing trend in DA concentration compared to the control group **(Figure 6)**. More interestingly, the NE levels in both antibiotic control groups and *C. difficile* challenged group were significantly reduced in the hippocampus (p < 0.05 and p < 0.001 respectively). This result suggests hippocampal DA release in response to *C. difficile* infection is distinctively different compared to striatum and prefrontal cortex, where dopaminergic circuits predominate.

**Figure 6.**
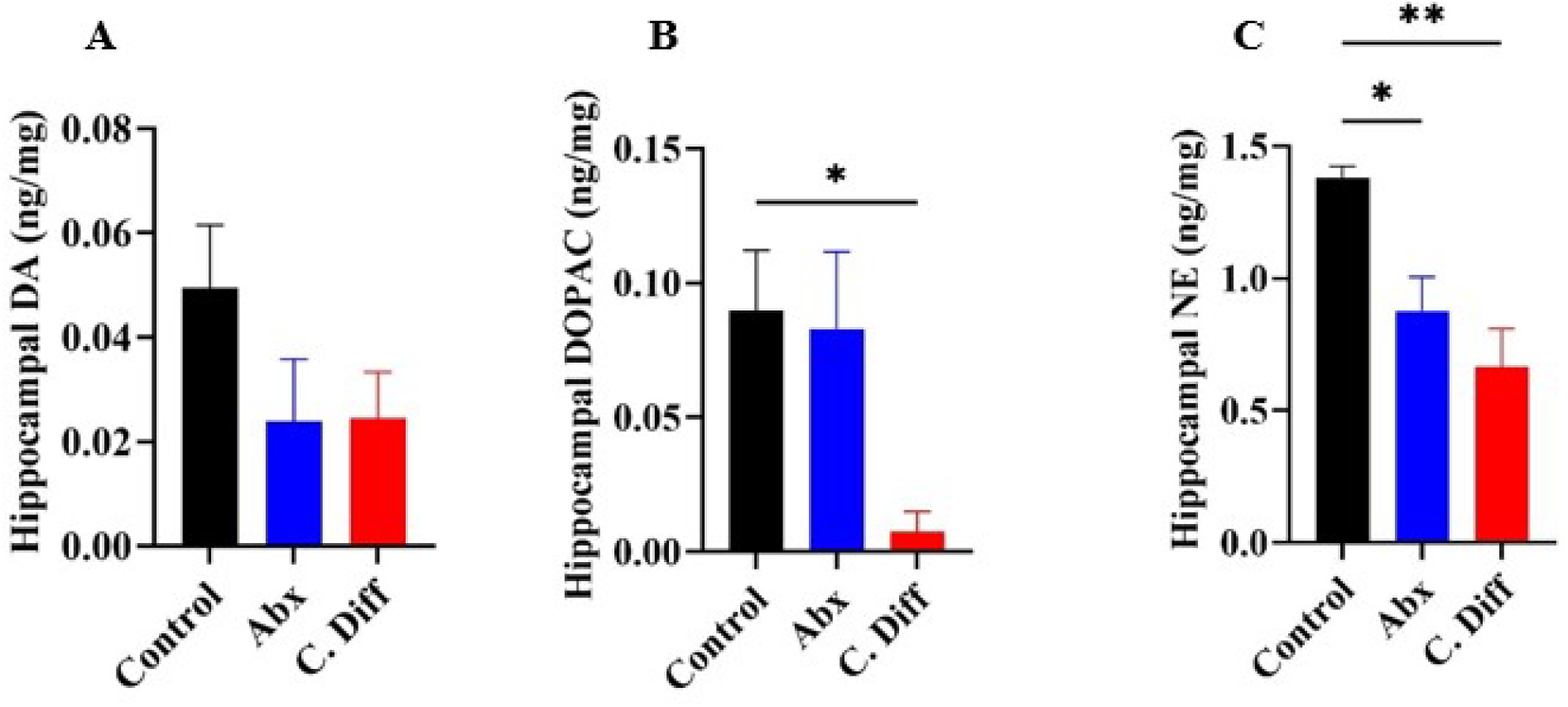
Antibiotic treatment and antibiotic-induced C. difficile infection decreased NE levels in the hippocampus: Three-four weeks old C57BL/6 mice are treated with an oral antibiotic cocktail or PBS and an intraperitoneal clindamycin injection or PBS for inducing gut dysbiosis and then challenged with 10^4^ *C. difficile* spores or PBS. Hippocampal catecholamines- **(A)** dopamine (DA), **(B)** its metabolite 3, 4-dihydroxyphenylacetic acid (DOPAC), and **(C)** norepinephrine (NE) levels in Control, antibiotic control (Abx), and *C. difficile* challenge (C. Diff) groups at 2-day post-inoculation were quantitated by HPLC-ED method. (HVA levels were below the detection limit in all groups) **p< 0.01, *p < 0.05.

### Antibiotic treatment, and antibiotic induced C. difficile infection differentially altered DA levels in substantia nigra

Substantia nigra is one of the major locations where neuronal DA is synthesized to be released to brain areas, such as striatum and prefrontal cortex through dopaminergic projections. Although not statistically significant, a decreasing trend in concentrations of DA and its degradation products were observed in the substantia nigra of *C. difficile* challenged mice compared to negative control **(Figure 7)**. In contrast, the concentrations of DA and DOPAC were significantly reduced in *C. difficile* challenged mice compared to the antibiotic control group (Abx) (p < 0.05). The same trend was also observed in HVA concentration. Moreover, the NE levels did not differ significantly across the treatment groups. The result suggests a reduced DA synthesis in substantia nigra in *C. difficile* infected animals (C.Diff group) but not in antibiotic (alone) treated animals (Abx group) compared to control, which will be further discussed in the following section.

**Figure 7.**
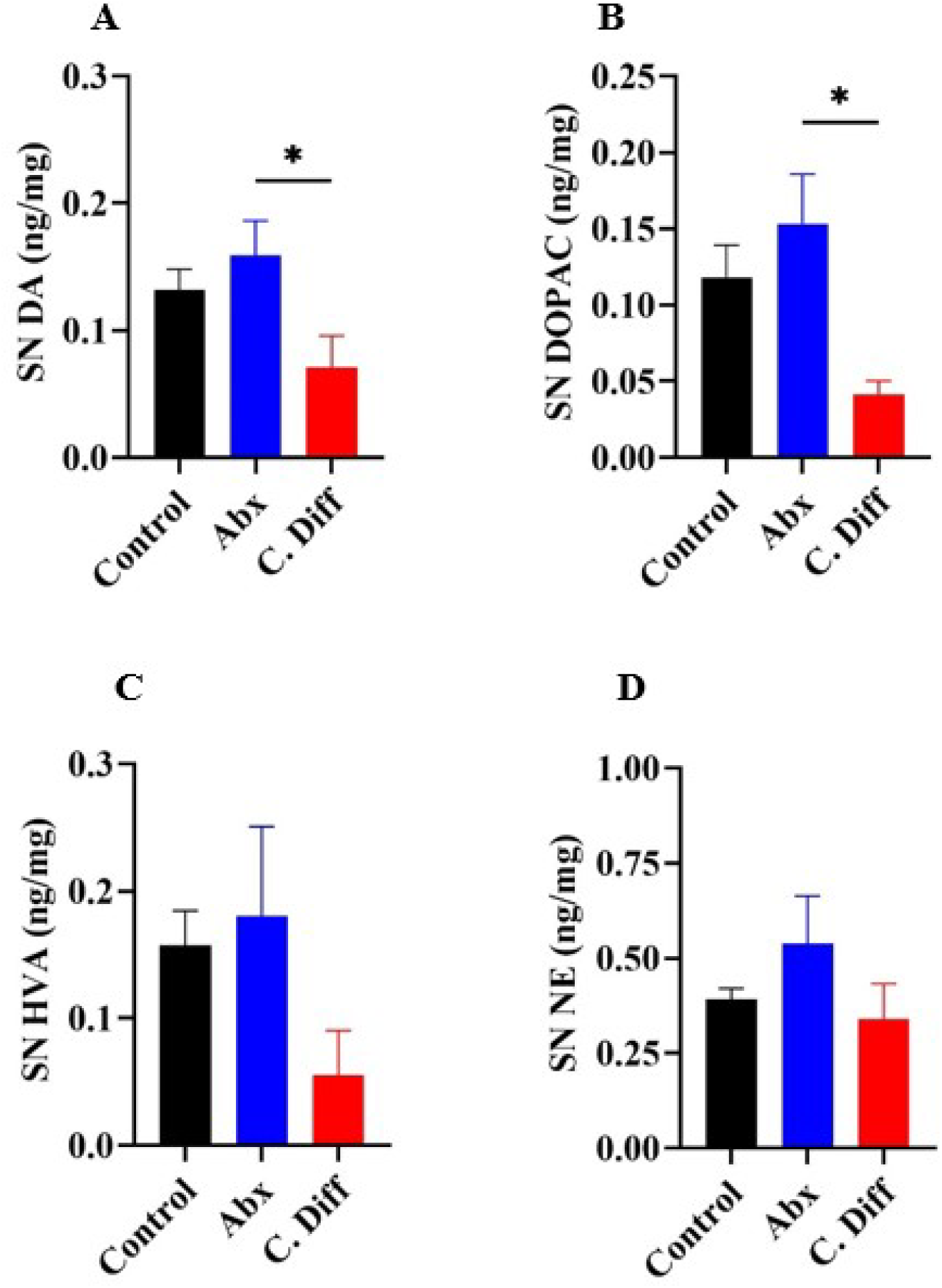
Antibiotic treatment and antibiotic-induced C. difficile infection differentially altered DA levels in substantia nigra: Three-four weeks old C57BL/6 mice are treated with an oral antibiotic cocktail or PBS and an intraperitoneal clindamycin injection or PBS for inducing gut dysbiosis and then challenged with 10^4^ *C. difficile* spores or PBS. Substantia Nigra **(SN)** catecholamines- **(A)** dopamine (DA), **(B)** its metabolite 3,4-dihydroxyphenylacetic acid (DOPAC), **(C)** homovanillic acid (HVA), and **(D)** norepinephrine (NE) levels in Control, antibiotic control (Abx), and *C. difficile* challenge (C. Diff) groups at 2-day post-inoculation were quantitated by HPLC-ED method. *p < 0.05.

## Discussion

Gastrointestinal disorders are one of the most common comorbidities reported in patients with ASD.^40–42^ Of particular note, gut dysbiosis, overgrowth of *C. difficile* in the gut, and gut microbiota-associated alterations in central neurotransmission have been implicated in ASD, where the dopaminergic axis plays an important role in the disease pathogenesis.^43^ In addition, a significant association has been demonstrated between antibiotic therapy and resultant alterations in the gut microbiome (gut dysbiosis)-which predispose *C. difficile* overgrowth in gut in early infancy contributing to ASD in children, although causation has not been proven.^44^ At this point, a specific underlying mechanism that connects gut dysbiosis to ASD-associated changes in the brain has not been identified.

In children and adults with ASD, gastrointestinal symptoms are a common problem frequently associated with dysbiosis and a higher incidence of *C. difficile* colonization^45, 46^. The severity of ASD has been strongly correlated with the severity gastrointestinal symptoms in autistic individuals^46^. Strikingly, fecal microbiome therapy (FMT), a radical treatment strategy against *C. difficile* infection in humans, is also found to be effective in mitigating ASD symptoms in both adults and children^47^. Additionally, transplantation of the fecal microbiome from ASD patients induced autism-like behavior in mice^48^. Moreover, vancomycin, a primary line of antibiotics used to treat *C. difficile* infection in humans, reduces autistic symptoms in affected individuals^49^. In contrast, fluoroquinolones, a group of antibiotic that predispose to the development of *C. difficile* infection in humans, aggravate ASD symptoms in autistic patients^49^. Thus, it appears that the factors associated with the predisposition, manifestation, and suppression of autistic symptoms coincide with that of *C. difficile* infection in affected individuals. Therefore, we hypothesized that gut-dysbiosis and dysbiosis-induced *C. difficile* overgrowth and subsequent production of p-cresol dysregulate the dopaminergic axis in the brain, which may have implications in neurodevelopmental disorders such as ASD.

As expected, the *in vitro* p-cresol production experiment showed that *C. difficile* UK1 strain efficiently converts the substrate p-HPA to p-cresol **(Figure 1)**. Production of a very high amount of p-cresol is an unusual property of *C. difficil*e compared to other gut bacteria.^50^ *C. difficile* ferments tyrosine and its metabolites such as p-HPA to synthesize p-cresol in the gut. p-Cresol is a bacteriostatic compound that impairs the membrane integrity of Gram-negative bacteria, which provides a competitive advantage to *C. difficile* over other bacteria, including *Escherichia* and *Bacteroides sp.*^51^ Similarly, the results from our infection trial indicated a significant increase in p-cresol concentration in the cecal contents of *C. difficile* infected mice compared to controls (p < 0.05) **(Figure 2D)**. It is known that p-cresol is readily absorbed into the bloodstream, as indicated by a significant increase in serum p-cresol levels in *C. difficile* infected animals compared to controls (p < 0.05). Although not statistically significant, an increased cecal p-cresol level was observed in the antibiotic control group (Abx). Previous reports, including our studies have already established that the antibiotic treatment (oral antibiotic cocktail and intraperitoneal clindamycin, in this case) causes significant dysbiosis in the murine gut, causing overgrowth of unfavorable bacterial communities that produce toxic metabolites, including p-cresol.^52, 53^ As expected, a similar trend was also observed in serum p-cresol levels of the antibiotic control mice (Abx group). These observations indicate that antibiotic-induced *C. difficile* infection increases serum levels of p-cresol, which may have significant extraintestinal, organ specific or systemic effects in the infected patient. Since p-cresol can cross the blood-brain barrier, it is reasonable to expect an impact on the structural and functional homeostasis of the central nervous system.

The enzymatic assay performed on the serum samples of *C. difficile* infected and control mice **(Figure 3B)** demonstrated a significant reduction in the DBH activity coinciding with the increased serum p-cresol levels. The brain is the primary source of the serum levels of DBH, and either serum or plasma DBH activity has been considered as one of the reliable measures of brain DBH activity.^36, 37^ It has been previously established that p-cresol is a potent inhibitor of this enzyme.^14–16^ DBH catalyzes the conversion of DA to NE in dopaminergic and noradrenergic neurons, and inhibition of DBH can increase the DA levels in the neurons and synapses with a concurrent reduction in the NE levels.^14–16^ The neurochemical profile of striatum and prefrontal cortex of the *C. difficile* challenged mice showed increased DA levels and a lower NE levels in these brain areas, suggesting that these alterations could be solely or partially a result of p-cresol mediated inhibition of DBH enzyme. However, further investigations are required to rule out the possibilities of reduced expression of DBH genes, and other pathways that affect neurotransmitter release in response to *C. difficile* infection.

The most notable observation made in this study is the increased DA in the striatum and pre-frontal cortex **(Figures 4 & 5)** with unchanged or decreased NE levels in mice with *C. difficile* infection (increased DA to NE ratio). These observations corroborate a recently published study which demonstrated increased DA levels in striatum and associated ASD like behavior in mice intravenously administered with p-cresol.^11^ DA release in striatum and prefrontal cortex is generally associated with reward response, motivation, pleasure, and addiction.^54^ In contrast, an increase in DA levels in these brain regions in response to persistent painful condition (dysbiosis and CDI associated typhlocolitis in this case) is unusual, as pain is a significant factor that ought to be considered in a *C. difficile* infection study. Generally, DA is considered as anti-nociceptive and persistent pain is often associated with a decreased DA levels in brain as opposed to acute pain, such as a pin prick.^55–58^ Therefore, a significantly increased DA and its degradation products observed in the striatum and pre-frontal cortex could be potentially attributed to a pathway that directly alters DA metabolism, such as p-cresol mediated inhibition of DBH. Increased DA (or its degradation products) to norepinephrine ratio as seen in both striatum and pre-frontal cortex of the *C. difficile* infected mice has been implicated in disorders of DBH activity, such as low DBH levels due to mutations in DBH coding genes.^37^ Therefore, *C. difficile* cresol mediated inhibition of DBH could be a potential reason for this phenomenon.

In contrast, the hippocampal neurochemical profile revealed an unaltered or decreasing trend of DA and DA degradation products in *C. difficile* infected animals **(Figure 6)**. Similar results have also been reported in a recent study, where the mice were administered intravenous p-cresol.^11^ This may suggest the dominance of the pain and memory-associated non-dopaminergic pathways in the hippocampus in response to *C. difficile* infection or high serum p-cresol. Interestingly, a significant decrease in NE is noticed in hippocampus, suggesting a reduced production of the NE in the nor-adrenergic neurons projecting to the hippocampus^59^, presumably due to p-cresol mediated inhibition of DBH. This enables exploration into β noradrenergic-mediated hippocampal-basolateral amygdalar control of social recognition memory (SRM) consolidation, which is believed to disordered in individuals with ASD and therefore fail to establish social relationship and social groups.^60^ In addition, we examined the catecholamine profile in the one of the two major dopamine producing areas in the brain that projects to the striatum, i.e., substantia nigra **(Figure 7)**. In addition to an unaltered NE level, the concentrations of DA, DOPAC and HVA appears to follow a decreasing trend, suggesting a potential feedback inhibition of DA production in response to accumulation of DA in striatum^61^. This is also in line with the hypothesis of altered DA to NE conversion in the brain in response *C. difficile* p-cresol mediated inhibition of DBH.

One of the major results from this study shows that *C. difficile* infection induces an increase in serum p-cresol, along with a concurrent increase in DA levels in the striatum, which has direct implications in the pathogenesis and aggravation of ASD-like behavior in autistic patients. A recent study by Pasucci *et al* demonstrated that intravenous p-cresol injection aggravates autistic behaviors in BTBR mice, a murine genetic model of ASD.^11^ Similar results are also demonstrated following oral p-cresol administration in non-mutant mice by Bermudaz-Martin *et al* ^13^. In Pasucci’s study, a single short-term intravenous p-cresol administration aggravated autistic behavior in mice, and most notably, increased the striatal DA levels. Interestingly, the serum p-cresol assay results and striatal neurochemical profile from *C. difficile* infected mice corroborates Pasucci *et al*’s observation. This study also suggested potential involvement of p-cresol mediated DBH enzyme inhibition to explain this observation. Similarly, Bermudaz-Martin *et al*’s reported impaired excitability of dopaminergic neurons and precipitation of autistic behavior in C57BL/6 mice continuously administered with oral p-cresol in drinking water. In contrast, Bermudaz-Martin *et al* attributed p-cresol mediated behavioral and neuronal alterations to remodeling of gut microbiota (gut dysbiosis) mediated by p-cresol, by demonstrating similar changes in mice transplanted with fecal material from p-cresol administered animals. However, Pasucci’s results indirectly nullify this theory by demonstrating similar behavioral and neurochemical alterations in short term, intravenous administration of p-cresol in BTBR, which entirely bypasses the involvement of gut and leaves minimal room for dysbiosis or microbial remodeling. Thus, an alternate but direct mechanistic pathway, such as inhibition of DBH, as proposed by Pasucci *et al* and our current study could be a plausible explanation for p-cresol induced alterations in dopaminergic neurotransmission. p-Cresol-producing bacteria, such as *C. difficile,* are extremely resistant to high gut cresol levels. Moreover, p-cresol strongly inhibits the growth of other bacteria in the gut providing a selective advantage for p-cresol-producing bacteria to overgrow.^50, 51^ This may result in further enhanced p-cresol production in the gut, and p-cresol mediated neurologic alterations in mice transplanted with such gut content, which more reasonably explains Bermudaz-Martin’s observation. Regardless, Bermudaz-Martin’s experiment compliments Pasucci *et al*’s study in several ways, as more prolonged nature of p-cresol administration in their study resulted in DA receptor insensitivity. It has been previously reported that prolonged, high DA levels and increase dopaminergic stimulation and impair DA receptor sensitivity in post synaptic neurons^62, 63^, supporting the results described in the current study and Pasucci *et al*’s report. Thus, the published studies, together with our current observations, suggest a potential direct link between *C. difficile* infection, gut and serum p-cresol levels, DBH activity, alterations in dopaminergic activity in brain that have direct implications in the pathogenesis of ASD. Therefore, the results from this study suggest a potential link between antibiotic induced *C. difficile* infection, bacterial p-cresol production, reduced DBH activity, and an altered dopaminergic axis, supporting the previous reports of p-cresol mediated ASD-like behavioral alterations in rodents.

Autism spectrum disorder is a group of heterogenous neurodevelopmental anomalies where genetic and environmental factors play complex roles, most of which are poorly understood. Our current study adds to the growing evidence of the ‘dopamine hypothesis of ASD’, which is at the same time, can be considered as a limitation of this investigation. Although dopaminergic system appears to be pivotal in the manifestation of classic autistic behavioral anomalies i.e., social deficits and stereotypic behavior, several other neurotransmitters and molecular pathways contribute to the pathogenesis of ASD. Another limitation of this study is that p-cresol may not be the only neurotoxic compound produced during dysbiosis and *C. difficile* infection in humans, and there could be several other unknown pathways implicated in reduced DBH activity and altered catecholamine metabolism in the brain. However, this study provides a logical association between gut dysbiosis, a known gut microbial toxin (p-cresol), and a potential mechanistic pathway of deregulated neurotransmission relevant to ASD. However, the scope of microbial mediated dopaminergic neurotransmission may not be restricted to ASD alone, but also applicable to other neurologic disorders such as Parkinson’s disease, where the dopaminergic neuronal pathology is implicated. For example, *C. difficile* has been recently shown to be associated with an increased short-term Parkinson’s risk, re-enforcing this concept.^64^

In summary, the current study demonstrated that antibiotic induced *C. difficile* infection causes significant alterations in the dopaminergic axis in mice. In addition, *C. difficile* infection significantly increased circulating p-cresol levels and reduced DBH activity in infected animals. Thus, the published studies, together with our current observations, suggest a potential direct link between *C. difficile* infection, gut and serum p-cresol levels, DBH activity, and alterations in dopaminergic activity in the brain that have direct implications in the pathogenesis of ASD. However, additional mechanistic, and behavioral studies are required to definitively confirm this hypothesis.

## Materials and methods

### Bacterial culture, media, and in vitro p-cresol production assay

The human epidemic *C. difficile* strain, UK1 (ribotype 027), was used in this study. *C. difficile* was cultured on two pre-reduced media: 1) BHI broth supplemented with yeast, 2) BHI broth supplemented with yeast and p-HPA, in anaerobic conditions (0% oxygen, 5% hydrogen, 5% CO2, and 90% nitrogen at 37°C) and incubated in an anaerobic workstation (AS-580, Anaerobe Systems, Morgan Hill, CA, USA). Primary *C. difficile* cultures were grown on BHI broth for 13 hours and then were diluted 1/100 into pre-equilibrated media with and without 0.3% p-HPA. After 24 hours, culture supernatant was collected after centrifugation at 5,000 x g at 4°C for 15 minutes, followed by filter sterilization. Supernatants were immediately aliquoted and saved at − 80 °C until analysis. All experiments were run in triplicates using pre-reduced media in an anaerobic environment, with every experiment being repeated twice. p-Cresol and p-HPA concentrations in the culture supernatant were estimated using HPLC method as per standard protocol.^51^

### Mouse model for antibiotic-induced C. difficile infection

All animal experiments were performed as per the protocols approved by ISU Institutional Animal Care and Use Committee (IACUC-20-091). Fifty-four C57BL/6 mice (3 to 4 weeks old) with an equal male-female ratio were purchased and housed in groups of 2 per cage with sterile food, water, and bedding. Mice were randomly assigned to each of the three groups: Control, antibiotic control (Abx), and antibiotic induced *C. difficile* infection (C. Diff) groups. After 4 days of acclimatization, a mixture of antibiotics and an intraperitoneal injection of clindamycin was administered to Abx and C. Diff groups, and the C. Diff group mice were orally gavaged with 1.4 x 10^4^ *C. difficile* spores.^34^ The animals were euthanized at approximately 48 hours post-inoculation, and the cecal contents and intestinal tissue samples, serum, and brain sections were collected for further processing.

### Histopathology

The cecum and colon tissues were fixed in 10% formalin and then embedded in paraffin. A 5μm thickness of tissue sections was made and stained with hematoxylin and eosin. A standardized scoring system was followed to assess CDI-associated histology injuries in intestinal tissues, and scores were evaluated across the groups based on epithelial damage, mucosal edema, and neutrophil infiltration.^65^ A board-certified pathologist (sample ID-blinded) performed microscopic analysis using an Olympus BX53 microscope (Olympus Optical Company, Tokyo, Japan). Scores were combined into a cumulative histologic score for each mouse ranging from 0 to 9.

### HPLC-ECD for tissue catecholamine analysis

Mouse brain regions including striatum, pre-frontal cortex, substantia nigra and hippocampus were micro-dissected with mouse brain matrix (ASI Instruments, RBM-1000S), preserved immediately in an antioxidant mixture containing 0.2 M perchloric acid, 0.1% NA_2_S_2_O_5_, 0.05% Na-EDTA, and isoproterenol (internal standard) as described previously and directly stored in dry ice.^66^ The samples were homogenized, sonicated, and filtered in 0.2-micron filter tubes. 20 µL samples were injected into High-Performance Liquid Chromatography – Electrochemical Detector (HPLC-ECD) with pump (Thermo-Scientific ISO-3100SD-BM), autosampler (Thermo-Scientific WPS-3000 TSL) and CoulArray 5600A-ECD detection system operated at 23 °C connected to Agilent Eclipse Plus (3.5 µm, 100A) C18 HPLC column (150×4.60 mm). Tissue neurochemical analytes were eluted in MD-TM MP mobile phase (cat. no. 701332, Thermo Fisher Scientific) under isocratic conditions at 0.6 ml/min for 21 min. DA (DA) and its major metabolites, Norepinephrine (N.E.), 3,4-dihydroxyphenylacetic acid (DOPAC), and homovanillic acid (HVA), were analyzed using CoulArray Data Station (v. 3.10) and quantified using a standard curve. Tissue neurochemical concentrations were normalized using wet tissue weight (mg) and corrected using the internal standard extraction coefficient.

### Quantification of p-cresol

Blood was collected by cardiac puncture. Serum was separated following clotting and centrifugation at 2500 ×g for 15 min. The collected serum was aliquoted as 40 µL into multiple microcentrifuge tubes and immediately temporarily stored in dry ice and further long-term storage at − 80 °C until analysis. The gut contents were collected and snap-frozen using liquid nitrogen and stored at − 80 °C until quantitative analysis via mass spectrometry.

#### Reagents

4-ethyl phenol, p-cresol (pC), salicylic acid (S.A.) were purchased from Millipore-Sigma, and p-cresol sulfate (pCS) and p-cresol glucuronide (pCG) were purchased from United States Biological, (Salem, MA). All the solvents were LC-MS grade and purchased from Fisher Chemical unless otherwise mentioned.

#### Instrumentation

Conjugated cresols were analyzed via an Agilent 6470 triple quadrupole (QQQ) mass spectrometer with electrospray ionization (ESI) equipped with an Agilent 1290 Infinity II UHPLC (Agilent Inc, MA, USA). Conjugated cresols were separated with the Agilent C18 column (2.1 x 100 mm, 1.8 µm) at 40°C. The mobile phase consisted of A: 5mM ammonium acetate + 0.005% acetic acid and B: Acetonitrile with a 0.400 ml/min flow rate. The solvent gradient was maintained at 0 min, 0% B; 0–0.25 min, 0–100% B; 0.5–15 min, 100-0% B; 15–20 min, with a 3-minute pre-equilibration.

#### Quantification

Analysis was performed in negative mode with the nozzle voltage set at 2000 V. Nitrogen was used as both nebulizer, desolvation, and sheath gas. The nebulizer pressure was set at 40 psi with a capillary voltage of 2500V. Desolvation gas was delivered at 12 L/min with temperature was heated to 350°C. The sheath gas flow was 10 L/min at a temperature of 350°C. High-purity nitrogen was used as a collision gas for collision induced dissociation. Multiple reaction monitoring (MRM) was used to detect cresol conjugates, the precursor, product ions, and collision energies are as follows: pCG precursor m/z 283 product m/z 107, 18 eV; pCS precursor m/z 187 product m/z 107, 18 eV; S.A. precursor m/z 137 product m/z 93, 25 eV.

Free p-cresol was analyzed with an Agilent 5975C quadrupole mass spectrometer equipped with 7890A gas chromatograph. The capillary column used was HP-88 (60m x 0.25mm x 0.2µm). An injection volume of 2 μL was used with the inlet operating in splitless mode. The oven GC program used an initial temperature of 40°C for 1 min, then increased to 230°C at a 20°C/min rate, with a final hold of 7.5 min. Inlet and MSD transfer line temperatures were held at 250°C. Mass detection was conducted under standard settings with a mass detection range set to m/z 40– 600. Peak identification was performed using Qualitative Analysis (version 10.0) software and the NIST Mass Spectral Search Program (version 2.3) with the NIST17 and Wiley 11 GC-MS spectral library. Peak detection and integration were accomplished using Agilent MassHunter Quantitative Analysis (version 10.0) software to monitor m/z 107 and m/z 108.

#### Sample preparation for conjugated p-cresol

The extraction protocol was adapted from Y. Morinaga et al. and C. Shu et al. with modifications as described here. ^6, 67^ The analysis of p-cresol conjugates was completed starting with 25-100µL sera samples or 150-500 mg gut contents previously archived in microcentrifuge tubes. Samples were kept on ice throughout the extraction. 4-ethyl phenol and salicylic acid were used as internal standards. These two internal control samples were spiked into the samples as follows: 4-ethyl phenol, 10 µL of 0.25mg/mL in ethyl acetate (2.5 µg); salicylic acid, 10µL of 0.05mg/mL in 50% methanol (0.5 µg). These internal standards were also spiked into 200 µL of 0.9% NaCl spiked with internal standards was used as a negative control. Three volumes (300µL/100mg-sample) of extraction solvent mixture of acetonitrile and water (2:1 v/v) were mixed with the sample, and after a quick vortex, this was incubated on ice for 10 minutes. The samples were sonicated for 10 min followed by a 10 min vortex. The supernatant was recovered after 7 min centrifugation at 13,000g. Pellets were saved in the freezer for free p-cresol extraction. 200 µL of the supernatant was filtered through a spin filter (catalog # UFC30LG25, Merck Millipore). Samples were subjected to LC-MS/MS analysis, 5 µL of which were directly injected into the LC-MS (QQQ).^67^

#### Sample preparation for free p-cresol

Analysis of free p-cresols was done with a continuation of the above extract for conjugate cresols. The supernatants were recombined with their respective pellets in the microcentrifuge tubes. Two volumes (200µL/100mg-sample) of 0.9% NaCl were added to the mix, followed by the addition of a solvent mix of 1:1 hexane: ethyl acetate (200µL). The mixture was sonicated for 10 min. After a 10 min vortexing, this was centrifuged for 7 min to obtain the phase separation. The top layer, which contains free cresols, was carefully transferred into a GCMS vial.

#### Preparation of calibration curves

Standards were made for conjugates in 2:1 acetonitrile:water 0.1-0.0005 mg/mL, and for p-cresol in ethyl acetate 0.75-0.0015mg/mL. 10µL of these standards were added in to 200µL 0.9% NaCl. Internal standards were also added and extracted by following the same extraction procedure for the samples.

### DBH activity assay

*Chemicals and reagents:* All assay chemicals were purchased from Millipore-Sigma unless otherwise stated. All solvents were LC-MS grade and purchased from Fisher Chemical.

#### Sample preparation and DBH assay reaction

Serum samples (40 µl) were prepared and assayed as previously described by Punchaichira et al., 2018, with the addition of LC-MS quantification as described below ^68–70^. Briefly, 40 μl of the serum samples were added to a cocktail of 1.0 mM tyramine HCl, 10 mM fumarate, 0.1 mg/ml catalase, 4 mM ascorbate, 2 μM copper sulphate, 30 mM NEM and 1.0 mM pargyline in 125 mM sodium acetate pH 5.2. The mixture was incubated for 1 h at 37 °C followed by addition of 50 μl of 25 mM EDTA for stopping the reaction.

#### Instrumentation

Equipment included Agilent Technologies 1100 Series HPLC system coupled to both a UV-Vis capable diode array detector (DAD) and an Agilent Technologies Mass Selective Trap SL detector equipped with a SphereClone 5 µm ODS (2) 80 A, L.C. Column 250 x 4.6 mm (Phenomenex) held at 40°C, and an electrospray ion source operated in positive mode. Running solvents consisted of A: 0.1% acetic acid in water and B: 0.1% acetic acid in acetonitrile. The flow rate was 0.700 mL/min with isocratic conditions of 35% B. The ion source drying gas was nitrogen which was set to flow at 12 mL/min at 350°C, the nebulizing gas was set to 30psi. The mass scan range was m/z 75-300 with a max accumulation time of 300 ms.

#### Quantification of enzyme activity

To quantify DBH activity, 10µL of each prepared serum DBH assay sample was injected for LC-MS analysis. The amount of octopamine converted from tyramine was quantified by monitoring the MS/MS product ion m/z 119 fragmented from the [M+H]^+^ ion (m/z 154) of octopamine at 0.7 fragmentation amplitude. An octopamine standard curve was generated in assay buffer ranging from 0-2 nmoles-octopamine/ sample. Quant analysis for 6300 Series Ion Trap LC/MS Version 1.8 (Bruker Daltonik, Bremen, Germany) software was used for the LC-MS/MS data analysis.

### Statistical analysis

Statistical analysis was performed using GraphPad Prism 9 (GraphPad Software, San Diego, CA) with p < 0.05 considered statistically significant. All results were expressed as means ± standard errors of the means (SEM) unless otherwise indicated. The differences between the experimental groups were compared using the analysis of variance (ANOVA). The differences between the two groups were analyzed using an unpaired Student’s t-test or Mann-Whitney test.

### Availability of data

The data supporting the findings of this study are available from the corresponding author SM on request.

